# SARS-CoV-2 infection of human neurons requires endosomal cell entry and can be blocked by inhibitors of host phosphoinositol-5 kinase

**DOI:** 10.1101/2022.09.14.508057

**Authors:** Pinja Kettunen, Angelina Lesnikova, Noora Räsänen, Ravi Ojha, Leena Palmunen, Markku Laakso, Šárka Lehtonen, Johanna Kuusisto, Olli Pietiläinen, Olli P. Vapalahti, Jari Kostinaho, Taisia Rolova, Giuseppe Balistreri

## Abstract

COVID-19 is a disease caused by coronavirus SARS-CoV-2. In addition to respiratory illness, COVID-19 patients exhibit neurological symptoms that can last from weeks to months (long COVID). It is unclear whether these neurological manifestations are due to infection of brain cells. We found that a small fraction of cortical neurons, but not astrocytes, were naturally susceptible to SARS-CoV-2. Based on the inhibitory effect of blocking antibodies, the infection seemed to depend on the receptor angiotensin-converting enzyme 2 (ACE2), which was expressed at very low levels. Although only a limited number of neurons was infectable, the infection was productive, as demonstrated by the presence of double-stranded RNA in the cytoplasm (the hallmark of viral replication), abundant synthesis of viral late genes localized throughout the neuronal cell, and an increase in viral RNA in the culture medium within the first 48 h of infection (viral release). The productive entry of SARS-CoV-2 requires the fusion of the viral and cellular membranes, which results in the delivery of viral genome into the cytoplasm of the target cell. The fusion is triggered by proteolytic cleavage of the viral surface protein spike, which can occur at the plasma membrane or from endo/lysosomes. Using specific combinations of small-molecule inhibitors, we found that SARS-CoV-2 infection of human neurons was insensitive to nafamostat and camostat, which inhibit cellular serine proteases found on the cell surface, including TMPRSS2. In contrast, the infection was blocked by apilimod, an inhibitor of phosphatidyl-inositol 5 kinase (PIK5K) that regulates endosomal maturation.

**Importance:** COVID-19 is a disease caused by coronavirus SARS-CoV-2. Millions of patients display neurological symptoms, including headache, impairment of memory, seizures and encephalopathy, as well as anatomical abnormalities such as changes in brain morphology. Whether these symptoms are linked to brain infection is not clear. The mechanism of the virus entry into neurons has also not been characterized. Here we investigated SARS-CoV-2 infection using a human iPSC-derived neural cell model and found that a small fraction of cortical neurons was naturally susceptible to infection. The infection depended on the ACE2 receptor and was productive. We also found that the virus used the late endosomal/lysosomal pathway for cell entry and that the infection could be blocked by apilimod, an inhibitor of the cellular phosphatidyl-inositol 5 kinase.

## Introduction

A variety of neurological symptoms have been observed in millions of COVID-19 patients which has led to a hypothesis that SARS-CoV-2 could infect brain cells. Such symptoms include fatigue, headache, impairment of concentration and memory (‘brain fog’), seizures, and encephalopathy (1). Structural changes in the brain anatomy have also been observed. A magnetic resonance imaging study of 785 participants found reductions in grey matter thickness and global brain volume in combination with changes in tissue contrast and tissue damage markers in certain brain areas (2). Post-mortem analysis of deceased COVID-19 patients has indicated sporadic presence of viral components in neurons, glial, and endothelial cells in different regions of the brain including the olfactory bulb (OB), which connects the olfactory sensory neurons of the nasal epithelium to the central nervous system (CNS) via a dense network of nerves (3–8). In *in vitro* cell culture models, SARS-CoV-2 can infect neurons derived from human embryonic stem cells (ESCs) and iPSCs in both two-dimensional (monolayers) and three-dimensional models (e.g., organoids) (7, 9–15). Some studies also reported infection in iPSC/hESC-derived astrocytes (8, 13, 15).

Bona fide neurotropic viruses such as rabies, poliovirus, or tick-borne encephalitis virus, cause severe neuronal infection that spreads to large areas of the brain with paralysing or lethal consequences (16–19). The potential of SARS-CoV-2 to infect very limited areas of the brain, and the possibility that this non-lethal infection could be transient, could explain some of the neurological manifestations in patients that suffer long COVID.

How SARS-CoV-2 enters brain cells is not clear. Also, whether the virus can spread from the initially infected neurons is debated, with studies showing both productive (10, 14) and non-productive infection (9, 12, 20).

Analysis of the SARS-CoV-2 infection in cell lines and primary respiratory epithelial cell models indicates that the virus has at least two possible entry routes: i) endocytosis and fusion from lysosomes or ii) direct fusion at the plasma membrane (21–23). Both mechanisms require a mildly acidic environment (pH <6.8) and the activation of the viral surface protein spike (S) by cellular proteases such as cathepsin-L (in lysosomes) or serine proteases such as TMPRSS2 (at the plasma membrane). Depending on the cellular availability of these proteases, infection can occur within lysosomes or at the cell surface. Inhibitors of endosome maturation (e.g., PIK5K inhibitors) block virus infection from endo-/lysosomes (21). Inhibitors of serine proteases (e.g., nafamostat and camostat) block infection from the plasma membrane. Here, we use authentic SARS-COV-2 to investigate the infection route and the spreading potential of the virus in 2D-cultured human iPSC-derived neurons, astrocytes and neuron-astrocyte co-cultures.

## Results and discussion

### Characterization of human iPSC-derived neurons and astrocytes

To study the viral entry mechanisms in human brain cells, we set up a human iPSC-derived neuron-astrocyte co-culture system in a 96-well plate format.

Firstly, we confirmed neuronal identity of the cells by positive staining for neuronal markers microtubule-associated protein (MAP2) and tubulin 3 (Fig. 1A & B). Furthermore, most of the neurons displayed a nuclear expression of Cux1, a marker for upper cortical layer neurons (layers II-III) (Fig. 1C) and lacked CTIP2, which is a marker for lower cortical layers V and VI (data not shown). In addition, most of the neurons showed robust staining with vesicular glutamate transporter 1 (Vglut1) (Fig. 1D). Together, these data imply that our cultures consist mainly of excitatory glutamatergic neurons of upper cortical layer (II-III) identity. However, some neurons displayed positive staining for GABA (Fig. 1B), which suggests that the cultures also contain small subsets of inhibitory GABAergic interneurons. The identity of iPSC-derived human astrocytes, obtained by a different induction protocol described in the methods, was confirmed by staining with astrocyte markers glial fibrillary acidic protein (GFAP) (Fig. 1E), S100β (Fig. 1F) and aquaporin 4 (AQP4) (Fig. 1G). The expression of GFAP and S100β mRNAs was further confirmed by qRT-PCR (Fig. 1H & I).

**Figure 1.**
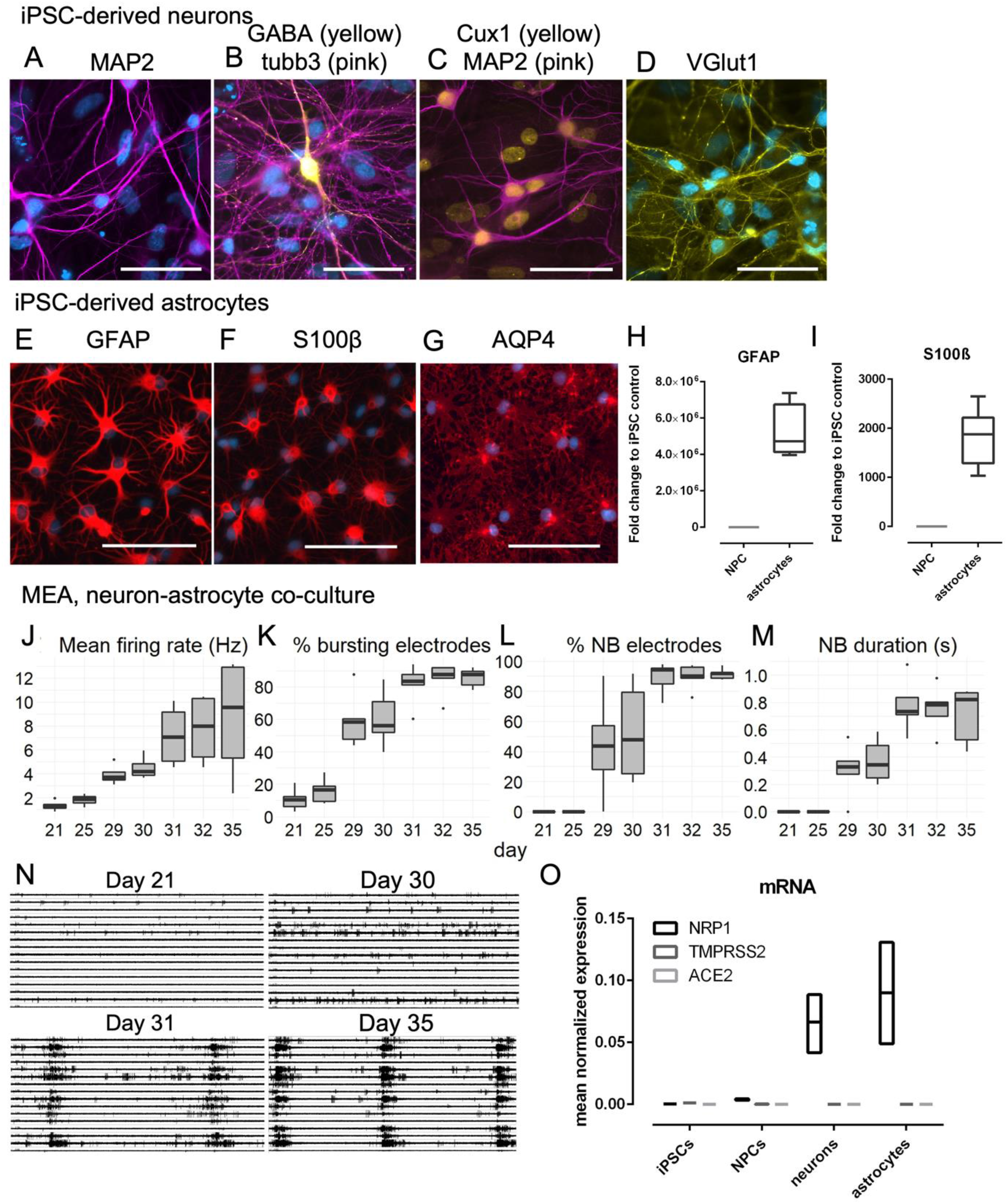
Characterization of hiPSC-derived neurons and astrocytes. A-D. Immunocytochemical staining of hiPSC-derived NGN2-neurons with MAP2, tubb3, GABA, Cux1 and Vglut1. Scale bar: 50 μm. N=5 cell lines. E-G. Immunocytochemical staining of iPSC-derived astrocytes with GFAP, S100β and AQP4. Scale bar: 100 μm. N=4 cell lines. H-I. qRT-PCR of GFAP and S100β expression in hiPSC-derived astrocytes. N=4 cell lines. J-M. Quantification of the mean firing rate (Hz), percentage of electrodes partaking in bursts, percentage of electrodes partaking in network bursts, and network burst duration (s). N=3 cell lines. N. Representative images of MEA recordings from neuron-astrocyte co-cultures at days 21, 30, 31 and 35. O. qRT-PCR of viral receptor expression in hiPSCs, NPCs, neurons, and astrocytes. N=2-4 cell lines.

Secondly, we used a microelectrode array (MEA) to estimate the maturity and functionality of the neuron-astrocyte co-cultures. The experiment showed that our neuron-astrocyte co-cultures develop an electrically active network capable of both single-electrode (Figure 1J-K) and network bursting (Fig. 1L). Cultures started developing electric bursting activity after three weeks of maturation (Fig. K) and network bursting appeared one week later (day 29, Fig. 1L). By day 31, most of the recording electrodes (>80%) participated in both single-electrode bursting and network bursting (Fig. K-L). Network burst duration increased until day 31 (duration 0.7s, Fig. 1 M). Representative images of MEA recordings from all 16 electrodes at days 21, 30, 31 and 35 are shown in Fig. 1N (10-second interval).

Finally, we used qRT-PCR to assess the expression of cell surface structures known to be important for SARS-CoV-2 infection in the respiratory tract. The expression of ACE2 and TMPRSS2 mRNA in iPSC-derived cortical neurons and astrocytes was detectable but very low (Fig. 1O). Both neurons and astrocytes expressed detectable levels of the entry co-factor neuropilin 1 (NRP1) (Fig. 1O), a protein that controls axonal development and has recently been implicated in SARS-CoV-2 infection (24, 25).

In summary, the neuron-astrocyte model appears to have electric activity typical of mature neurons, correct cell markers, and cells endogenously express viral entry factors.

### Infection of iPSC-derived neural cultures by SARS-CoV-2 is mainly dependent on ACE2 receptor and does not spread efficiently

To assess susceptibility of iPSC-derived human neural cultures to SARS-CoV-2, we infected 30-day-old neuron-astrocyte co-cultures with the ancestral SARS-CoV-2 Wuhan strain and analyzed samples by immunofluorescence analysis of viral protein expression at various time points. A representative image of an infected well at 48 hours post infection (hpi) stained with DNA dye Hoechst 33342 (nuclear marker), neuronal-specific marker microtubule-associated protein 2 (MAP2), and SARS-CoV-2 nucleocapsid protein (N) is given in Fig. 2A, with an enlarged area shown in Fig. 2B. The viral N protein is distributed both in the cell body, the soma, and throughout neurites (dendrites), (Fig. 2B, arrow heads).

**Figure 2.**
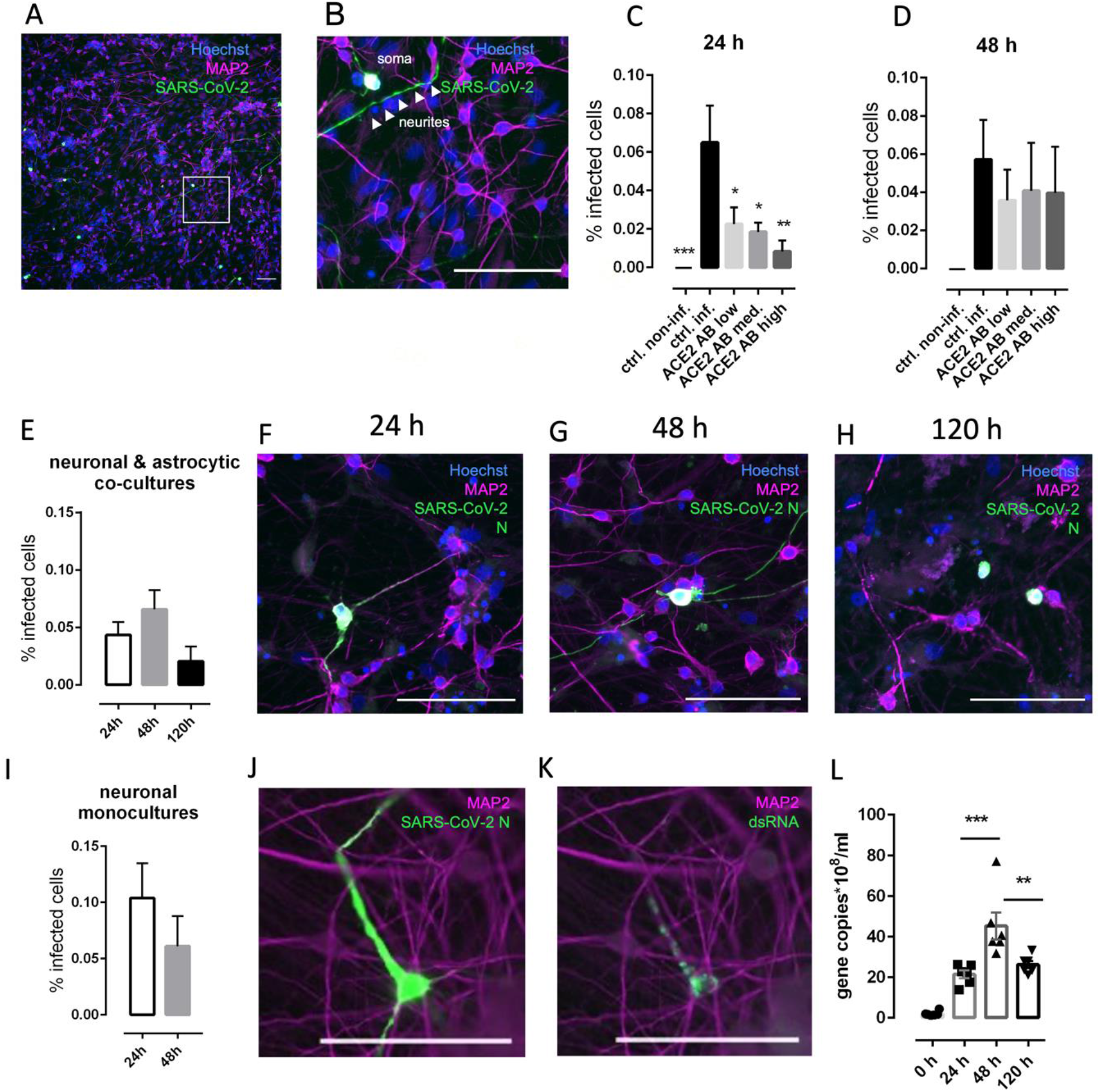
Infection of hiPSC-derived neural cultures by SARS-CoV-2 is mainly dependent on ACE2 receptor and does not spread efficiently. A. Representative image of SARS-CoV-2 N protein staining in the infected hiPSC-derived co-cultures of neurons and astrocytes (48 hpi). B. Enlarged area from the box in Fig. 2A. C. Infection of neurons is blocked by anti-ACE2 antibody in a dose-dependent manner at 24 hpi. N=6 samples per group. D. Infection of neurons is not blocked by anti-ACE2 antibody when the virus remains in the medium for 48 h. N=6 samples per group. E. Percentage of the infected cells in co-cultures consisting of neurons and astrocytes (all the infected cells were identified as neurons based on their MAP2 expression) does not change at different time points tested. N=6 samples per group. F, G, H. Representative images of neurons co-cultured with astrocytes, infected with SARS-CoV-2 and stained with N protein collected at 24, 48 and 120 hpi. L. Percentage of the infected cells in neuronal monocultures does not change at different time points and is similar to the infection level in neurons cultured with astrocytes. N=6 samples per group. H. Staining of an infected cell with MAP2, anti-N and (I) anti-dsRNA antibody that demonstrate their colocalization. J. qRT-PCR of the cell medium at different time points post infection. N=6 samples per group. All scale bars are 100 μm. Columns and bars represent mean ± SEM, respectively. Data were analyzed by (C) ordinary one-way ANOVA followed by Dunnett’s multiple comparisons test, (D, E) ordinary one-way ANOVA, (L) unpaired t test, (J) ordinary one-way ANOVA followed by Tukey’s multiple comparisons test. *p<0.05, **p<0.005, ***p<0.0005.

ACE2 receptor is the primary receptor used by SARS-CoV-2 to enter cells (26). Some of the earliest studies on SARS-CoV-2 have challenged the possibility for SARS-CoV-2 infection in the central nervous system due to the low ACE2 mRNA levels found in the human brain (27, 28). Since then, other studies have found robust ACE2 protein expression in human neurons (7, 29), with Song et al. further reporting that application of anti-ACE2 antibody prior to infection could block SARS-CoV-2 in human brain organoids. We found that SARS-CoV-2 is able to infect neurons but not astrocytes. To test whether SARS-CoV-2 infection of human iPSC-derived neurons is dependent on ACE2, we treated the cells with different concentrations of anti-ACE2 antibody (2 ug/mL, 5 ug/mL, 20 ug/mL) 1 h prior to infection with SARS-CoV-2 (1.5 MOI). At 24 h, the application of anti-ACE2 Ab significantly blocked the infection in neurons in a dose-dependent manner (Fig. 2C). At 48 h, application of anti-ACE2 Ab was less effective (Fig. 2D), perhaps due to the consumption of the antibody by the cells. Overall, these data confirm previous findings that the neuronal SARS-CoV-2 infection is at least partly ACE2-dependent, which is similar to SARS-CoV-2 infection in other cell types (30, 31).

Following the infection with SARS-CoV-2 at a multiplicity of infection (MOI) of 1.5, we found a low level of infection (around 0.05%) at all time points we analyzed (24, 48 and 120 h) (Fig. 2E). All of the N-positive cells demonstrated robust staining for neuronal-specific marker MAP2, suggesting that all of the infected cells were neurons. No astrocytes (defined as MAP2 negative cells) were infected in the experiment. Our data support previous findings where neurons, but not astrocytes, are susceptible to SARS-CoV-2 infection in human iPSC-derived cultures (9, 11, 12, 14). Some previous studies observed SARS-CoV-2 infection in astrocytes, too (13, 15). However, both studies that found SARS-CoV-2 infection of astrocytes have used a faster cell differentiation protocol with the astrocytes attaining a more immature morphology than the astrocytes used in our study. Since it has been shown that neural progenitor cells (NPCs) are susceptible to SARS-CoV-2 infection (7, 14), it is possible that the maturity of astrocytes may affect their infectability, with more immature astrocytes being more vulnerable to SARS-CoV-2 infection. We, therefore, focused on determining the infectious entry pathway of SARS-CoV-2 in neurons.

The level of the neuronal SARS-CoV-2 infection did not differ significantly between the analyzed time points (one-way ANOVA, Fig. 2E), which is in line with a previous study (12). It is also interesting that even though the infection stage of the N+ cells was not always the same at a specific time point, we could still observe emergence of a pattern. At 24 h, the infection was mostly localized in the neuronal soma, with proximal dendrites beginning to display a sign of infection (Fig. 2F). At 48 h, it was common to see fully infected cells, with all the neurites showing a robust positivity for SARS-CoV-2 N (Fig. 2G). At 120 h, all the cells that were found positive for N had their neurites retracted, which is a sign of a severely diseased state (Fig. 2H).

Since previous work by Wang et al. demonstrated that the presence of astrocytes exacerbates neuronal susceptibility to SARS-CoV-2 infection in human iPSC neuronal cells (13), we challenged iPSC-derived neuronal monocultures with a similar dose of SARS-CoV-2 that we used to infect neuron-astrocyte co-cultures. The level of infection in neuronal monocultures was comparable to the level of infection in neuron-astrocyte co-cultures, suggesting that astrocytes do not facilitate neuronal SARS-CoV-2 infection in these cultures (Fig. 2I).

Additionally, we confirmed colocalization of anti-N and anti-double stranded RNA (dsRNA) antibodies (Ab) in the infected samples, demonstrating presence of both viral protein and viral RNA material (Fig. 2J and 2K). To check whether the infection of neurons is productive, we carried out qRT-PCR analysis of the medium collected from the cells at 0, 24, 48 and 120 hpi. We observed that viral genome was released into the medium, with the maximum load detected at 48 hpi (Fig. 2L).

### Virus infection is blocked by inhibition of PIK5K but not serine proteases

To infect cells, SARS-CoV-2 surface protein spike (S) has to be cleaved by cellular proteases, which is followed by fusion of the virus with the membrane of the cell or its components. Previous studies reported that SARS-CoV-2 could infect human primary cells: 1) through endocytosis, spike activation by cathepsin-L and fusion of the virus with lysosomes, or 2) through activation of the spike by transmembrane serine protease 2 (TMPRSS2) and direct fusion with the plasma membrane (21, 22, 32). To investigate which route of infection is utilized by SARS-CoV-2 in human iPSC-derived neurons, we used drugs to block these pathways, alone or in combination. Apilimod blocks 1-phosphatidylinositol 3-phosphate 5-kinase (PIK5K) and therefore disrupts endosomal/lysosomal trafficking, which has previously been shown to block viral infections, including Ebola and SARS-CoV-2 (33–35). It eliminated SARS-CoV-2 infection in neurons at 24 h (Fig. 3A) and significantly reduced it at 48 h when applied 1 h prior to infection with SARS-CoV-2 at 1.5 MOI (Fig. 3B). Nafamostat, that inhibits serine proteases and prevents the virus from entering the cells directly from the plasma membrane, did not block the infection (Fig. 3A and 3B). A combination of both drug types had an effect similar to apilimod alone (Fig. 3 A and 3B).

**Figure 3.**
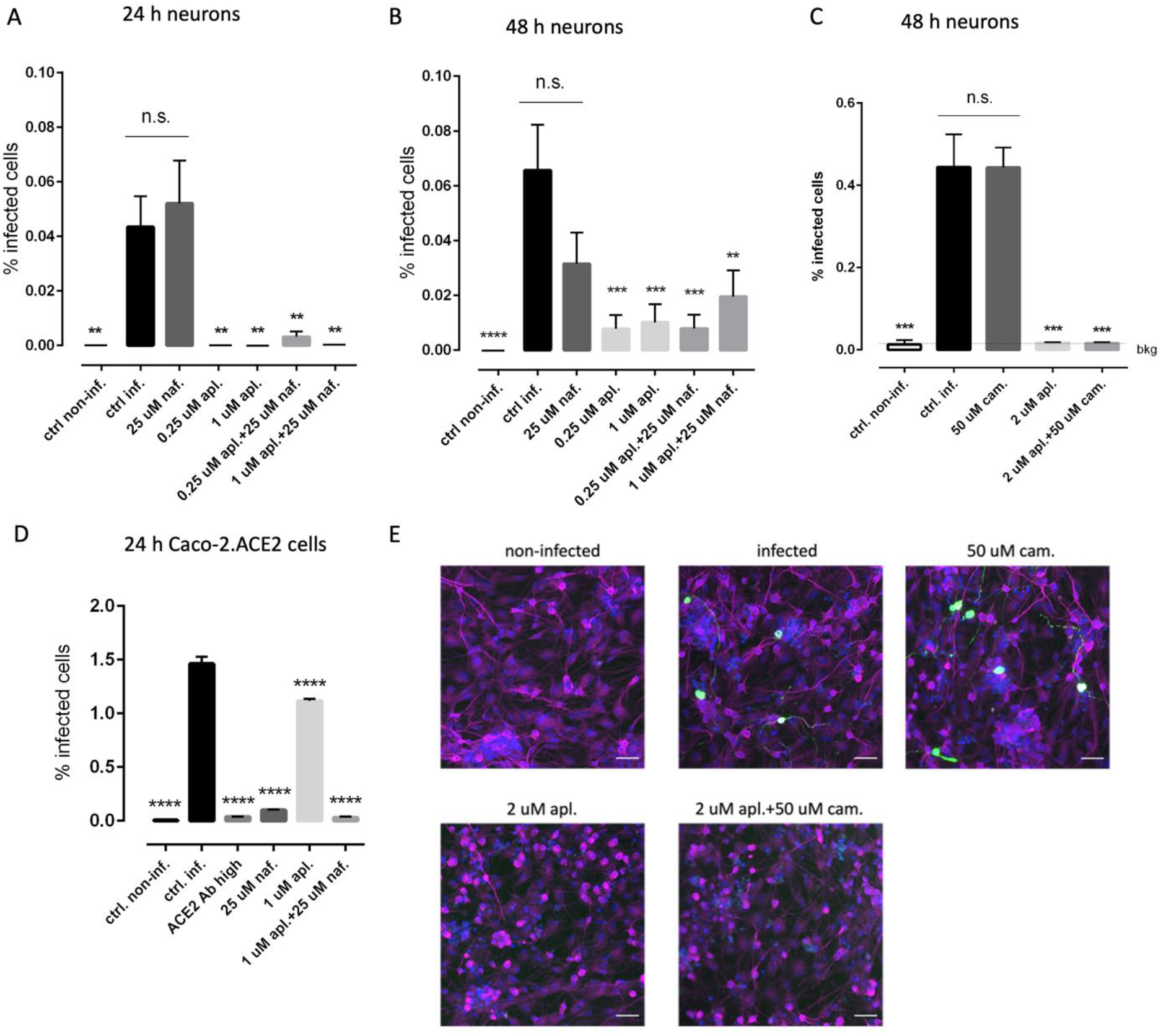
SARS-CoV-2 infection of hiPSC-derived neural cultures is blocked by inhibition of PIK5K but not serine proteases. A. Infection of neurons can be blocked by apilimod but not by nafamostat at 24 h at 1.5 MOI. N=6 samples per group. B. Infection of neurons can be blocked by apilimod but not by nafamostat at 48 h at 1.5 MOI. N=6 samples per group. C. Infection of neurons can be blocked by apilimod but not by camostat at 48 h at 15 MOI. N=3 samples per group. D. SARS-CoV-2 infection can be effectively blocked by nafamostat in Caco-2 cells expressing ACE2, where the major cell entry route for the virus is through the plasma membrane and not through late endosomes/lysosomes. N=3 samples per group. E. Representative images of SARS-CoV-2 infection (staining with N protein) in neural cultures at 48 h at 15 MOI. Blue – Hoechst 33342, magenta – MAP2, green – SARS-CoV-2 N. All scale bars are 100 μm. Columns and bars represent mean ± SEM, respectively. Data were analyzed by ordinary one-way ANOVA followed by Dunnett’s multiple comparisons test.

Since our initial level of infection in neurons was low, we decided to check whether an increase in the viral titer might increase the level of infection. Therefore, we infected the cells with SARS-CoV-2 at MOI of 15 and evaluated the infection at 48 hpi. While we observed around a 10-fold increase in the infection of the cells, the overall infection rate was still low (around 0.5%) (Fig. 3C). Camostat, another serine protease inhibitor analogous to nafamostat, had no effect on infectivity of SARS-CoV-2 in neuronal cells, while apilimod (alone or in combination with camostat) robustly blocked the infection.

To control for the overall effectiveness of the drugs, we used the Caco-2 cell line expressing ACE2 receptor. We treated the cells with similar concentrations of apilimod and nafamostat and infected the cells with SARS-CoV-2 at 2.5 MOI 1 h post treatment. While apilimod had only a small effect on the infection rates in Caco-2 cells, nafamostat rendered a remarkable decrease in the infection levels in Caco-2 cells, confirming previous data that cell surface serine protease inhibitors are capable of blocking SARS-CoV-2 entry in cells where this route is available for the virus (Fig. 3D). Since neuronal infection was blocked by an inhibitor of the host factor phosphatidyl-inositol 5 kinase but not by inhibitors of cell surface serine proteases, these data suggest that SARS-CoV-2 infection of iPSC-derived neurons preferentially occurs through the endosomal pathway and not through direct fusion with the plasma membrane preceded by TMPRSS2-mediated cleavage. Therefore, drugs that disrupt viral entry through the endosomal/lysosomal pathway could possibly be used in preventive care or soon after the exposure to the virus. However, we warn against hasty or incautious use of such drugs. In our *in vitro* experiments, apilimod negatively affected the morphology of neurons by causing neurite truncation (Fig. 3E). Thus, apilimod has served as a useful tool to evaluate the SARS-CoV-2 entry pathway, but it is an unlikely candidate for clinical trials unless it is carefully tested safety in preclinical studies *in vivo*.

## Conclusions

The current study has characterized SARS-CoV-2 infection in human iPSC-derived cortical neurons and provided evidence that neurons but not astrocytes get infected even at higher viral doses. The infection relies on ACE2 for entry and it is productive. When entering the neuronal cells, the virus preferentially uses the endosomal/lysosomal pathway. SARS-CoV-2 requires at least three factors to infect cells: 1) a receptor (e.g. ACE2), 2) a protease to activate the fusogenic activity of the spike (e.g. cathepsins in endo/lysosomes or TMPRSS2 at cell surface), and 3) a pH < 6.8 (23). While the average pH of human nasal mucosa is indeed around 6.6 (23), potentially allowing virus fusion at the PM, the pH of extracellular fluids in the brain is rather neutral, above 7.2 (36). It is therefore conceivable that the virus requires endocytosis and access to acidic endosomes to infect neurons. Drugs that inhibit any of these steps could potentially be used in preventive care or soon after the infection exposure to prevent or limit neuron infection, respectivelly.

Very low-level infection in the brain might not be easily traced, especially if not lethal. However, even low-level infection could lead to long-lasting negative consequences. Although the infection led to neuronal cell death within 120 h *in vitro*, we do not know how long an infected neuron could release viruses and survive *in vivo*. A deeper understanding of brain infection by SARS-CoV-2 could, on the one hand, help understand if there is a casual connection between the virus infection of brain cells and the neurological manifestations associated with Long COVID. On the other hand, a more detailed molecular characterization of the virus entry pathways and mechanisms of assembly and release are needed to develop treatments against COVID-19-associated neurological complications.

## Materials and methods

### Generation and culturing of human iPSCs

Punch skin biopsies were collected from Finnish healthy males after informed consent. The study has received acceptance from the Research Ethics Committee of the Northern Savo Hospital District (license no. 123.13.02.00/2016). Skin fibroblasts were expanded in fibroblast culture media (Iscove’s DMEM, 20% fetal bovine serum, 1% penicillin-streptomycin and 1% non-essential amino acids) as described previously (37) and transduced using CytoTune™ –iPS 2.0 Sendai Reprogramming kit (ThermoFisher Scientific) according to manufacturer’s instructions. Fibroblast culture medium was replaced with Essential 6 Medium (E6 supplement, ThermoFisher Scientific) supplemented with 100 ng/ml basic fibroblast growth factor (bFGF, Peprotech) at day 6 post-transduction. Reprogrammed iPSC colonies were selected based on morphology at three weeks post-transduction and expanded on Matrigel (growth-factor reduced, Corning) in Essential 8 (E8) Medium at 37°C / 5% CO_2_. Cultures were passaged with 0.5 mM EDTA approximately twice a week. The pluripotency of newly generated hiPS cells was verified by immunocytochemistry for stage-specific embryonic antigen 4 (SSEA-4), octamer-binding transcription factor 4 (Oct4), Nanog, and TRA-1-81 (Supplementary Figure 1). All used iPSC lines are listed in Supplementary Table 1.

### hiPSC-derived NGN2-neurons

hiPSC-derived neurons were generated according to the protocol adapted from Nehme et. al. 2018 (38). Briefly, hiPSCs were transduced with a lentivirus containing the NGN2-gene under tetracycline-inducible promotor (Tet-O-Ngn2-Puro) in combination with lentivirus carrying FUdeltaGW-rtTA construct (both plasmids from Addgene, lentivirus packing and concentration by Alstem) at MOI 10 for 1 h. The construct contains a puromycin-resistance gene, which allows for selection of neural precursor cells (NPCs). After transduction, the virus was removed, and the cells were cultured normally on Matrigel (growth-factor reduced, Merck) -coated 35 mm culture plates in E8-medium (Gibco) + 50 U/ml penicillin + 50 μg/ml streptomycin). NGN2-transduced iPSCs were expanded under normal hiPSC culture conditions and stock vials were frozen in 10% DMSO (Sigma), 10% fetal bovine serum (Biowest) in culture medium.

For neuronal differentiation, a vial of NGN2-transduced hiPSCs was thawed and passaged 1-3 times under normal culture conditions prior to use. On day 0, neuronal differentiation was initiated by adding 2 μg/ml doxycycline to E8 medium on a 60-70% confluent NGN2-iPSC plate. On day 1, medium was switched to N2-medium (DMEM/F12 without L-glutamine, 1%, Glutamax, 1% N2 (all from Gibco), 0.3% glucose) supplemented with 2 μg/ml doxycycline (BioGems) and dual SMAD inhibitors 0.1 μM LDN-193189 (Sigma), 10 μM SB-431542B (Sigma), 2 μM Xav939 (BioGems).

On day 2, developing NPCs were selected by adding 5 μg/ml puromycin (ThermoFisher Scientific). On day 3, puromycin was removed, dead cells were washed away with base medium and the cells were returned to N2-medium supplemented with 2 μg/ml doxycycline, 0.1 μM LDN-193189, 10 μM SB-431542B, 2 μM Xav939. On day 4, emerging neurons were plated with or without astrocytes on 0.9 - 13 mm coverslips or glass-bottom 96-well plates coated with 9-16 μg/cm^2^ poly-d-lysine and ∼1.5 μg/cm^2^ laminin (from mouse Engelbreth-Holm-Swarm (EHS) sarcoma; Sigma). Density was 50 000 cells / cm^2^/ cell type. Medium was switched to Neurobasal (Gibco) supplemented with 1% Glutamax (Gibco), 2% B27 without vitamin A (Gibco), 50 μM non-essential amino acids (Gibco), 0.3% glucose, and 10 ng/ml GDNF, BDNF and CNTF (Peprotech). Day 7: proliferation was inhibited with an overnight 10 μM floxuridine (Tocris) -treatment. The cells were maturated for 4-6 weeks with 50% medium changes three times a week.

### hiPSC-derived astrocytes

Astrocyte differentiation was initiated by growing confluent hiPSC plate in Neural Maturation Medium (Neurobasal, DMEM/12 without L-glutamine, 1.7% Glutamax, 50 μM non-essential amino acids, 0.5 mM sodium pyruvate, 0.5% N2, 1% B27 with vitamin A, 50 μM beta-mercaptoethanol, 2.5 μg/ml insulin, 50 U/ml penicillin, 50 μg/ml streptomycin) supplemented with dual SMAD inhibitors 10 μM SB-431542B and 200 μM LDN-193189 for 10-12 days. Resulting NPCs were split 1:2 by scraping and plated on 1.5 μ/cm^2^ laminin-coated 35 mm cell culture dishes. The NPCs were expanded for 2-4 days in NMM supplemented with 20 ng/ml bFGF. Then the cells were detached and moved to ultra-low attachment plates (Corning) in Astrodifferentiation medium (DMEM/F12 without L-glutamine, 1% Glutamax, 50 μM non-essential amino acids, 1% N2, 50 U/ml penicillin, 50 μg/ml streptomycin, 0.5 UI/ml heparin) supplemented with 10 ng/ml bFGF and EGF. Astrospheres were cultured for 6 months and cut manually when necessary. For experiments, astrospheres were dissociated with StemPro Accutase for 10 min, triturated into a single-cell suspension and plated on culture dishes. For characterization, astrocytes were plated on growth factor-reduced Matrigel (Corning) at density 50 000 cells/cm^2^ and maturated in Astrodifferentiation medium supplemented with 10 ng/ml BMP-4 & CNTF.

### Immunocytochemistry for cell type characterization

Cell cultures were fixed with 4% paraformaldehyde (PFA) and washed twice with PBS. Cells were permeabilized for 20 min with 0.25% Triton X-100 in PBS or left unpermeabilized (SSEA-4 and TRA-1-81). Unspecific binding was blocked with 5% normal goat serum in PBS (blocking buffer). Primary antibodies were diluted in blocking buffer and incubated overnight in 4°C. The next day, the samples were washed 3 × 10 min with PBS. Secondary antibodies were diluted 1:1000 in blocking buffer and incubated for 1 h at room temperature (RT). Samples were washed 3 × 10 min with PBS and stained with nuclear marker DAPI prior to mounting with Fluoromount. Characterization was done for six neuronal cell lines (N=6) and four astrocyte cell lines (N=4).

### Immunohistochemistry for virus-infected samples

The cells were fixed with 4% PFA in PBS at RT for 20 min. PFA was removed, and the cells were incubated in PBS at 4° C until the staining was performed. The virus was inactivated with ultraviolet radiation (5000 J/m^2^ dose) before removal of the samples from BSL-3. Before permeabilization, the cells were incubated in 50 mM ammonium chloride (NH_4_Cl) in PBS to quench free aldehyde groups remaining post fixation at RT for 20 min. Then the cells were permeabilized with 0.1% Triton-X in PBS and the nuclei were stained with 1:1000 Hoechst 33342 in Dulbecco medium containing 0.2% bovine serum albumin (BSA-Dulbecco) for 10 min. The cells were washed once with 0.2% BSA-Dulbecco and incubated in primary Ab at 4° C overnight. On the following day, the cells were washed twice with 0.2% BSA-Dulbecco and incubated in fluorescent dye-conjugated secondary antibodies for 1 hour at RT. After that, the cells were washed with 0.2% BSA-Dulbecco three times and 100 μl of PBS per well was added.

### Antibodies

Full list of antibodies used in the study is provided in Supplementary Table 2.

### Microelectrode array (MEA)

Electric activity of neuron astrocyte-cocultures was assessed using the Maestro Edge multi-well microelectrode array system (Axion). The cells were plated on 24-well Cytoview MEA plates (Axion) at density 60 000 neurons + 60 000 astrocytes per well and cultured for 3 weeks prior to starting the recordings. Each well contained 16 electrodes per well with 50 μm electrode diameter and 350 μm electrode spacing. The activity was recorded at 37°C / 5% CO_2_ for 10 min until day 35. The signal was sampled at frequency 12.5 Hz and filtered with digital low pass filter 3 kHz Kaiser Window and digital high pass filter 200 Hz IIR. The noise threshold was set at 5 standard deviations. Bursts were detected with the following inter spike interval (ISI) threshold settings: minimum number of spikes: 10; maximum interspike interval: 100 ms. Network bursts were detected with the following settings: minimum number of spikes: 90; maximum interspike interval: 20 ms; minimum percentage of participating electrodes: 33%. Characterization was done using five cell lines (N=5) and all values were calculated as a mean of three wells.

### qRT-PCR to assess RNA expression of astrocyte markers and cell surface receptors

Levels of ACE2, GFAP, S100β NRP1 and TMPRSS2 receptors in our hiPSC-derived neurons, astrocytes, and neuronal precursor cells (NPCs) were assessed with qRT-PCR. First, RNA was isolated from cultured neurons, astrocytes, NPCs and iPSCs using RNeasy mini kit (Qiagen) following manufacturer’s instructions. The concentration of RNA was measured using NanoDrop and 500 ng of RNA was converted into cDNA. First, 500 ng of RNA was diluted in water and mixed with Random hexamer primer (ThermoFisher Scientific). The samples were incubated 5 min at 65°C in C1000 Thermal Cycler (Bio-Rad). Then, a synthesis mixture (10 mM dNTP, ribonuclease inhibitor and Maxima reverse transcriptase in reaction buffer (ThermoFisher Scientific) was added to the samples and cDNA synthesis was run for 30 min in 50°C. Quantitative RT-PCR was run using Maxima probe/ROX qPCR master mix and the following TaqMan® primers: *ACE2* (HS01085333_m1), *GAPDH* (Hs99999905_m1), *GFAP* (Hs00909233_m1), *Neuropilin-1* (Hs00826128_m1), *S100β* (Hs00902901_m1), *TMPRSS2* (HS01122322_m1) (Thermo Fisher Scientific) on Bio-Rad CFX96 Real-Time System (Bio-Rad). The samples were run at 95°C for 10 min followed by 40 cycles of 95°C 15 s, 60°C 30 s, and 72°C 30 s. The results were normalized to human *GAPDH* expression using the Q-gene program (Equation 2) (39). qRT-PCR was repeated for tree cell lines each (N=3) with two exceptions: four (N=4) and two (N=4) astrocyte cell lines were used to assess ACE2 and NRP1 RNA expression, respectively.

### qRT-PCR to assess viral release

SARS-CoV-2 RNA was harvested from the cell medium at various time points post infection and stored in viral lysis buffer with RNA supplements (50 μl of sample medium in 560 μl of AVL buffer supplemented with 1% carrier RNA) (Qiagen). RNA was extracted using QIAamp Viral RNA Mini Kit (Qiagen). RNA concentration and quality were evaluated using NanoDrop 2000 Spectrophotometer (ThermoFischer Scientific). qRT-PCR was run in triplicates using reverse transcriptase- and template-negative controls and TaqMan Fast Virus 1-step MasterMix by Thermo Fischer Scientific (5 μl of the sample in 20 μl of total volume). Primers and probe used for the reaction were ordered from Metabion: RdRP-SARSr-F2: 5’-GTG ARA TGG TCA TGT GTG GCG G -3’ as the forward primer, RdRP-SARSr-R2: 5’ - CAR ATG TTA AAS ACA CTA TTA GCA TA - 3 as the reverse primer’, RdRP-SARSr-P2: 5’-6 - Fam - CAG GTG GAA CCT CAT CAG GAG ATG C -BHQ - 1 - 3’ as the probe. The qRT-PCR reaction was run with AHDiagnostics Agilent Technologies Stratagene Mx3005P using the following steps: reverse transcription for 5 min at 50°C, initial denaturation for 20 s at 95°C and two amplification steps at 95°C for 3 s and 60°C for 30 s (the amplifications steps were repeated for 40 cycles).

### SARS-CoV-2 virus

Wuhan strain of SARS-CoV-2 virus produced in Vero E6 cells was used in all the experiments.

### Anti-ACE2 antibody treatment

Different concentrations of anti-ACE2 Ab (low – 2 μg/ml, medium – 5 μg/ml, high – 20 μg/ml) were added to the cells 30 min. prior to the infection with the virus.

### Drug treatment

The cells were treated with 0.25 μM, 1 μM, 2 μM apilimod dimesylate (Tocris, ref. #7283, batch 1A/257560), 50 μM camostat mesylate (Tocris, ref. #3193, batch 2B/242261), 25 μM nafamostat mesylate (Tocris, ref. #3081, batch 6A/257562) or combinations of these drugs 30 min prior to the infection with the virus.

### Virus Infections

Experiments involving infection of cells with SARS-CoV-2 were performed in BSL-3 facility of the University of Helsinki under all required university permissions. Infection of cells was performed in Neurobasal media (Gibco) supplemented with 1% Glutamax, 2% B27 without vitamin A, 50 μM non-essential amino acids, 0.3% glucose, and 10 ng/ml GDNF, BDNF and CNTF. Before application to the cells, SARS-CoV-2 was pre-incubated in the neurobasal medium at 37° C for 30 min. After the infection, the cells were incubated at 37° C with 5% CO_2_ supplementation for 24, 48 or 120 h. For Caco-2 cell experiment, infection was carried out in Dulbecco’s Modified Eagle’s Medium (Sigma, D6546) supplemented with 4500 mg / L glucose, sodium pyruvate, sodium bicarbonate, 1% L-glutamine, 1% NEAA, 2% fetal bovine serum, penicillin and streptomycin.

### High-throughput imaging to detect the virus

High-throughput imaging was carried out using ImageXpress Nano microscope (Molecular Devices) at the Light Microscopy Unit of University of Helsinki. We used two Nikon objectives: 10x/0.3 Plan Fluor, WD 16 mm (pixel size 0.655 μm) and 20x/0.45 S Plan Fluor ELWD, WD 8.2-6.9 mm (pixel size 0.328 μm). Detailed information on the filter specifications for different channels can be found at https://wiki.helsinki.fi/display/LMU/MolecularDevices+Nano.

### Image Analysis

High-throughput image analysis was carried out using Cell Profiler 4 (40). Approximately 10000 cells per sample were analyzed. First, the nuclei were identified on the image channel dyed with Hoechst 33342 by providing a typical diameter of the objects using Otsu thresholding method. Then the objects identified as nuclei were expanded by a few pixels to approximately represent the borders of a cell. The channel with the MAP2/Tubulin-3 staining was overlaid with the expanded nuclei, and a threshold for an average signal intensity of MAP2/Tubulin-3+ cells was chosen. The expanded nuclei displaying an intensity of the MAP2/Tubulin-3+ staining channel above the chosen threshold were classified as neurons, while the cells with the intensity below were classified as astrocytes. Both classes of cells were overlaid with an image channel where staining for the virus N-protein was carried out. Cell bodies demonstrating intensity of staining above a defined threshold were counted as virus-positive cells. Percentage of cells positive for the virus N-protein from the total population of the cells of their cell type was plotted. For the control experiment in Caco-2 cells, the pipeline was similar, but no cell type-specific markers were used as the cell line was homogeneous.

### Statistical analysis

Statistical analysis of the data was performed in GraphPad Prism 6. ROUT test was performed before the analysis to identify outliers. Two-tailed unpaired Student’s t-test was performed when two conditions were compared. One-way ANOVA was used when more than two groups were analyzed. When the main effect was found statistically significant, post-hoc multiple comparisons tests were carried out. Differences were considered statistically significant when p<0.05. Data in figures are presented as mean ± SEM.

## Data availability

We are in the process of submitting the data underlying the current research into a public repository. DOIs of the original research data will be included into the manuscript text before the official publication.

## Acknowledgments

We thank Ida Hyötyläinen, Laila Kaskela, Eila Korhonen, Jenni Voutilainen for the work they have done on iPSC characterization. The study was supported by the Sigrid Juselius Foundation (J.K., Š.L.), The Academy of Finland (Grant 334525, JK), University of Helsinki Graduate Program in Microbiology and Biotechnology (R.O.), Academy of Finland Research Grants 335527 (G.B., A.L. and R.O.) and 336490, Jane and Aatos Erkko Foundation (O.V., Š.L.), Helsinki University Hospital funds TYH2021343 (O.V.), European Union’s Horizon Europe research and innovation program grant 101057553 (G.B., O.V.) and University of Helsinki Doctoral Programme Brain & Mind (P.K). The funders had no role in the study design, data collection and interpretation. High-throughput imaging was performed at Light Microscopy Unit of the University of Helsinki. The authors declare no competing interests.

## Supplementary material

**Sup. Fig. 1.**
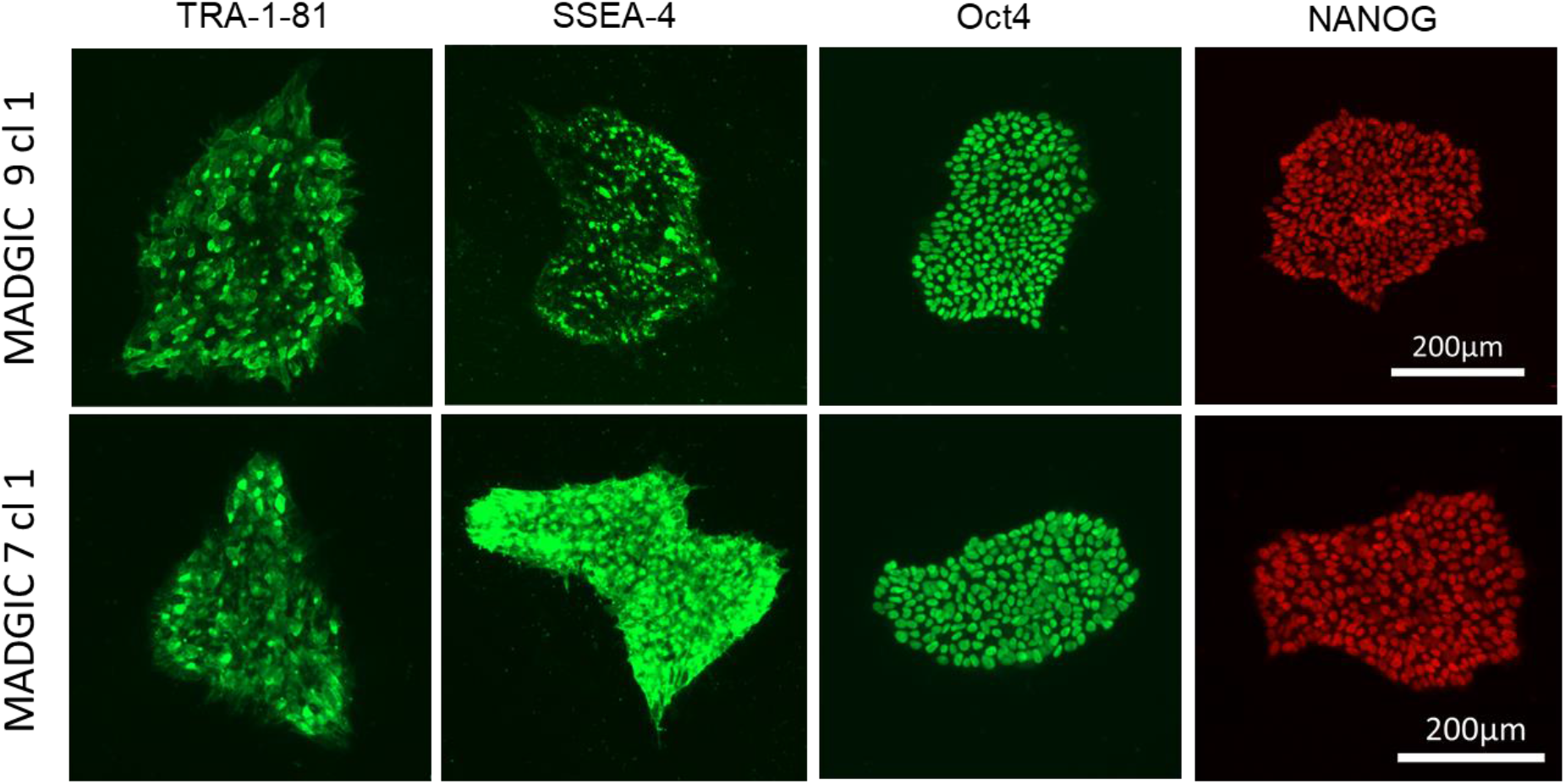
Immunocytochemical characterization of previously unpublished hiPSC lines (MADGIC 7cl1 & MADGIC 9cl1) with TRA-1-81, stage-specific embryonic antigen 4 (SSEA-4), octamer-binding transcription factor 4 (Oct4) and NANOG antibodies. Scale bar 200 μm.

**Sup. Fig. 2.**
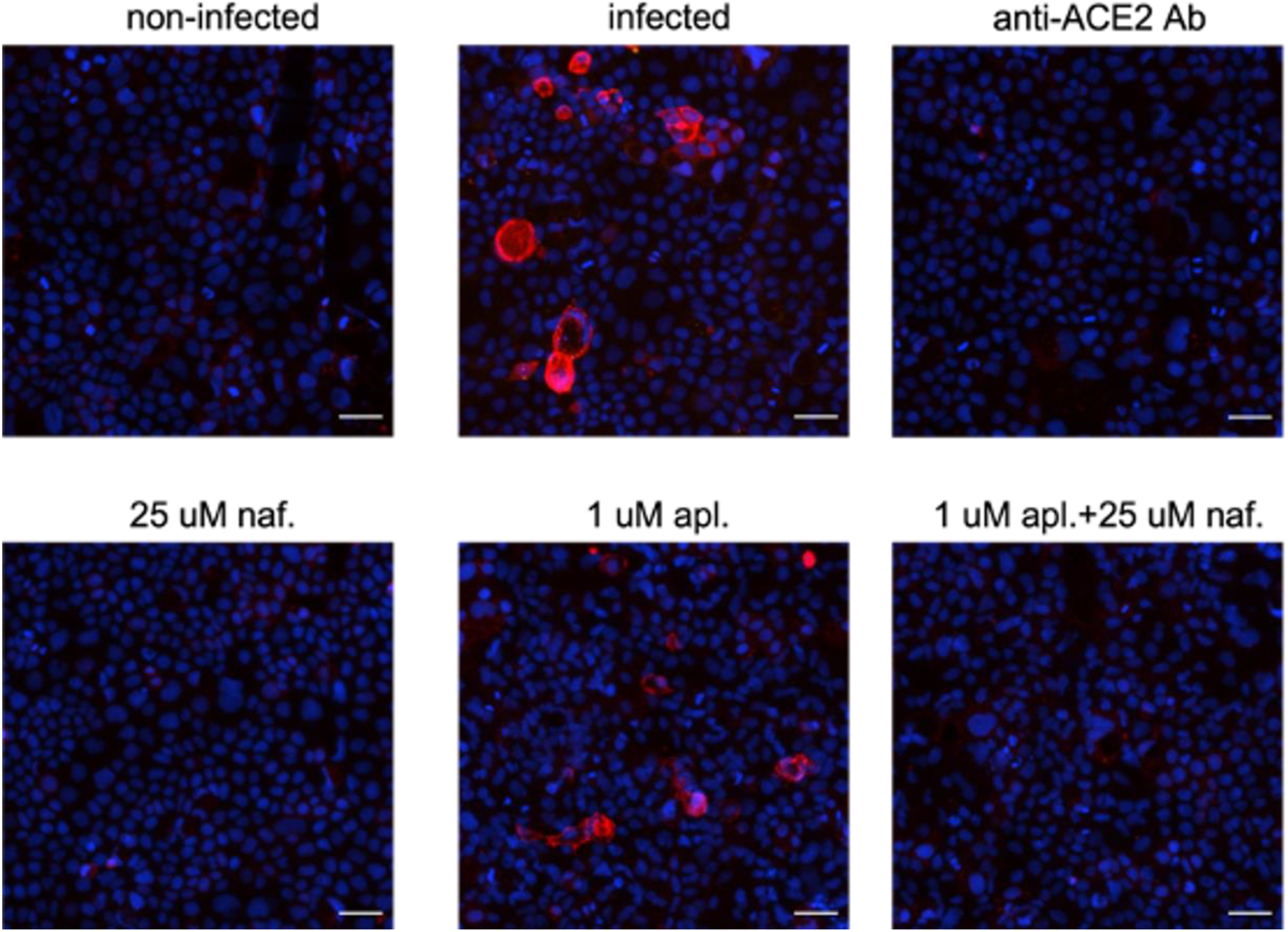
Representative images of SARS-CoV-2 infection (staining with N protein) in Caco-2.ACE2 cells at 24 hpi at 2.5 MOI. Blue – Hoechst 33342, red – SARS-CoV-2 N. Scale bar 100 μm.

**Sup. Table 1.** A list of the hiPS cell lines used in this study.

| ID | Gender | Age | Cell type | Published in |
| --- | --- | --- | --- | --- |
| MADGIC 1cl7 | male | 67 | Astrocyte | Jäntti et. al 2022 |
| MADGIC 4cl1 | male | 65 | Astrocyte,<br>neuron | Lehtonen et. al. 2018 |
| MADGIC 6cl1 | male | 63 | Neuron | Jäntti et. al 2022 |
| MADGIC 7cl1 | male | 66 | Neuron | Previously unpublished |
| MADGIC 8cl1 | male | 64 | Astrocyte,<br>neuron | Jäntti et. al 2022 |
| MADGIC 9cl1 | male | 58 | Neuron | Previously unpublished |
| MADGIC 12cl2 | male | 58 | Astrocyte,<br>neuron | Rolova et. al. 2020 |

**Sup. Table 2.** Primary and secondary antibodies used in the study.

| Antibody | Concentration<br>used | Manufacturer | Reference<br>number |
| --- | --- | --- | --- |
| ACE2 goat | 1:1000 | R&D Systems | AF933 |
| CTIP2 rat | 1:500 | Abcam | ab18465 |
| Cux1 mouse | 1:500 | Abcam | ab54583 |
| GABA rabbit | 1:500 | Sigma | A2052 |
| GFAP rabbit | 1:500 | Agilent | Z033429-2 |
| MAP2 chicken | 1:1000 | Abcam | ab92434 |
| Nanog | 1:100 | Thermo Fisher<br>Scientific | MA1-017 |
| Nucleocapsid<br>rabbit | 1:2000 | gift from Jussi Hepoioki | (38) |
| Oct4 | 1:400 | Millipore | MAB4401 |
| S100β rabbit | 1:500 | Abcam | ab52642 |
| SSEA4 | 1:400 | Millipore | MAB4304 |
| TRA-181 | 1:200 | Millipore | MAB4381 |
| Tubulin-3 mouse | 1:2000 | Biolegend | 801201 |
| Vglut1 rabbit | 1:500 | Sigma | V0389 |
| Goat anti-mouse<br>488 | 1:400 | Molecular Probes | A11001 |
| Goat anti-mouse<br>Alexa 633 | 1:500 | Invitrogen | A21052 |
| Goat anti-rabbit<br>Alexa 488 | 1:1000 | Invitrogen | A11008 |
| Goat anti-chicken<br>Alexa 568 | 1:1000 | ThermoFisher | AB_2534098 |
| Chicken anti-goat<br>Alexa 488 | 1:1000 | Invitrogen | A21467 |
| Donkey anti-rabbit Alexa 647 | 1:1000 | Invitrogen | A31573 |
| Donkey anti-mouse Alexa 555 | 1:1000 | Invitrogen | A32773 |

